# Individual variation in male pheromone production in *Xylocopa sonorina* correlates with size and gland color

**DOI:** 10.1101/2023.09.27.559843

**Authors:** Andrew J. Goffinet, Kathy Darragh, Nicholas Saleh, Madeleine M. Ostwald, Stephen L. Buchmann, Santiago R. Ramirez

## Abstract

Sex pheromones are species-specific chemical signals that facilitate the location, identification, and selection of mating partners. These pheromones can vary between individuals, and act as signals of mate quality. Here, we investigate the variation of male pheromones in the mesosomal glands of the large carpenter bee *Xylocopa sonorina*, within a Northern California population. We tested the hypothesis that morphological traits are correlated with the observed variation in chemical blend composition of these bees. We also conducted behavioral assays to test whether these male pheromones act as long-range attractants to conspecifics. We found that larger males with darker mesosomal glands have a higher pheromone amount in their glands. Our analysis also shows initial evidence that this pheromone blend serves as a long-range attractant to both males and females. We show that both male body size and sexual maturation are important factors influencing pheromone abundance, and that this pheromone blend acts as a long-range attractant. We hypothesize that this recorded variation in male pheromone could be important for female choice.

## INTRODUCTION

Detection of chemical signals is considered to be amongst the most ancient and widespread of sensory systems, playing a major role in both inter- and intraspecific communication (Ache and Young 2005; Amo and Bonadonna 2018). In insects, pheromones are interspecific chemical signals, which are important for a diverse range of behaviors including aggregation (Wertheim et al. 2005), trail recruitment, and social organization (Czaczkes et al. 2015), and mating (Wyatt 2014). The chemical signals themselves can be comprised of a single compound (Daimon et al. 2012) or a mix of multiple compounds (Andersen et al. 1988). A particularly well-studied group of chemical signals are those involved in mating, known as sex pheromones (Wyatt 2014).

Sex pheromones can serve a multitude of purposes during mating and mate choice. In some cases they are important for the initial step of mate attraction, acting over long ranges to attract individuals of the opposite sex (Itagaki and Conner 1988; Baker 1989). They are often species-specific signals, maintaining reproductive isolation by reducing hybridization between sympatric species (Daimon et al. 2012). Sex pheromones can also play key roles in intraspecific mate choice and/or signaling mate quality to the receiver, two roles that are not mutually exclusive (Juárez et al. 2015).Sex pheromones are thought to act as honest signals of mate quality, as diet can impact pheromone production (Darragh et al. 2019), with males fed a higher quality diet showing increased reproductive success (Rantala et al. 2003; Liedo et al. 2013). Therefore, while pheromones are often species-specific, they also show variation both between and within populations due to genetic drift, selection, or variation in diet quality or environmental conditions (Allison and Cardé 2016).

The large carpenter bees (Apidae: *Xylocopa*) are a cosmopolitan genus of wood nesting bees, which display a range of mating behaviors and male secondary sexual traits. Ancestrally, *Xylocopa* males defend resources to gain access to females, which is correlated with small or no mesosomal glands and a lack of sexual dimorphism (Leys and Hogendoorn 2008). However, in some subgenera, males defend non-resource based territories (Minckley 1994). These species tend to have sexually dimorphic males, with males exhibiting enlarged mesosomal glands, potentially some of the largest exocrine glands in insects (Minckley 1994). These glands release long-range sex pheromones to attract females and are thought to signal indirect benefits. An especially well-studied species in this group is *Xylocopa sonorina*, the Valley Carpenter Bee, found from California, Arizona, New Mexico, to Texas and into Sonora, Mexico, and introduced to Hawaii and various Pacific Islands. This species displays extreme sexual dimorphism, with all-black females and golden males with green eyes (Figure 1) (Marshall and Alcock 1981). In early spring, males leave their nests daily in the late afternoon, where they defend a non-resource based territory (Marshall and Alcock 1981; Alcock and Smith 1987). These are often referred to as a lek mating system (as in some birds) where one or more male carpenter bees simultaneously display to arriving females as potential mates (Marshall and Alcock 1981). Despite female interactions being relatively infrequent, the majority of these interactions do not result in copulation (Marshall and Alcock 1981; Alcock and Johnson 1990). Additionally, given that females must land to commence copulation, and that the males do not pursue unmated females, mate choice appears to be driven primarily by the female (Alcock and Smith 1987; Alcock and Johnson 1990).

**Figure 1:**
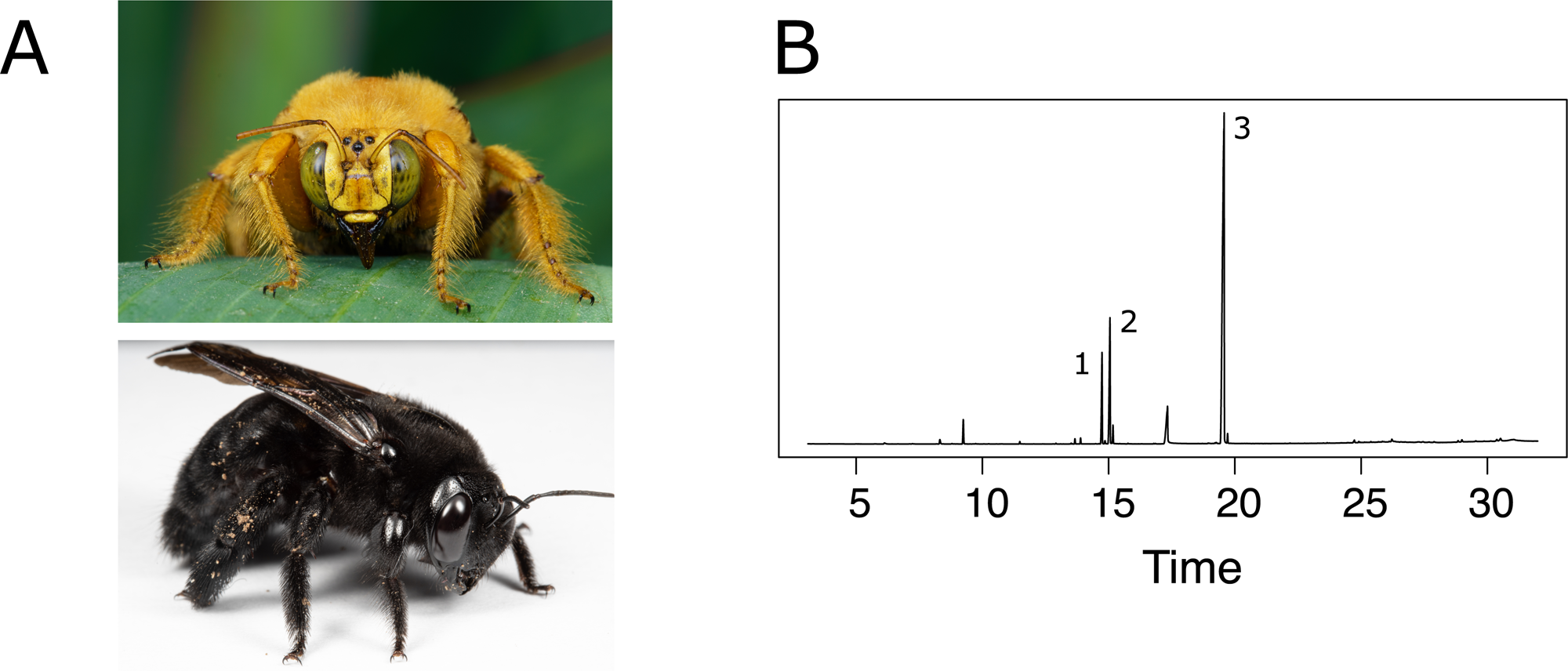
(a) Male *X. sonorina*. Photo credit: Sebastian Scofield. (b) Representative chromatogram of a male’s mesosomal gland extract, showcasing the three primary compounds and the range of minor constituents. The peak labeled as 1 is 3,7,11-methyl-2,7,10-dodecatrienal, the peak labeled as 2 is all trans-farnesal, and the peak labeled as 3 is all trans-geranylgeraniol.

Sex pheromones play an important role in male *X. sonorina* mating displays. Males choose a focal point, either a branch or a sprig of a non-flowering shrub or tree, which they scent-mark and hover above, either chasing away or flying in loops with conspecific males (Marshall and Alcock 1981; Alcock and Smith 1987). The pheromone released by males is easily detectable to human observers (Marshall and Alcock 1981), who perceive it as a distinctive rose-scent, with the primary constituent being a common compound found in rose volatiles (Dani et al. 2021). One hypothesis is that the similarity of this pheromone to floral scents could have evolved to exploit pre-existing sensory systems used to find flowers for nectar sources (Schiestl 2010). The pheromone itself is a blend of three primary compounds: all trans-geranylgeraniol, all trans-farnesal and 3,7,11-methyl-2,7,10-dodecatrienal in a 9:6:1 ratio (Andersen et al. 1988). In a previous field bioassay, the two most abundant components seem to attract females as successfully as wild males (Minckley et al. 1991). Producing the pheromone is thought to be costly as the mesosomal glands, which are lacking in females, are very large, occupying almost 20% of thoracic volume, otherwise occupied by the indirect flight muscles in other bee species (Ostwald et al. 2022). The quantity of male sex pheromone produced potentially indicates male quality, with females seeming to assess several males before making a decision (Marshall and Alcock 1981). Across *Xylocopa* species, enlarged glands are associated with long-distance mate attraction(Minckley et al. 1991); within a species, larger glands may similarly store greater quantities of pheromones and facilitate attraction of females at greater distances.

The mesosomal gland undergoes a distinct ontogenetic development process throughout the adult lifespan of *X. sonorina* males. In pre-sexual males, who overwinter as adults with siblings in their natal nests, the gland exists in a static, non-secretory phase. The following spring, the gland enters a secretory phase and pheromone production begins approximately a month before they begin to leave their nests to seek mating opportunities at lek sites (Buchmann pers. obs.). At some point during the mating season, the gland darkens in color as cuticular sclerotization replaces the pale-colored former secretory tissue cell layers with hard, setal-lined hollow tubules (Ostwald et al. 2022). This transition marks the ends of the gland’s secretory phase, with the gland now functioning as a storage organ for the accumulated pheromone blend (Buchmann pers. obs). In this way, gland color can indicate its developmental stage.

In this study, we investigate how the chemical profile of *Xylocopa sonorina* males varies within its population from Northern California. We analyze the mesosomal gland contents released by displaying males during their active season. We describe the pheromone blend found within the glands and identify factors correlated with pheromone variation, including body size and gland color, both of which may covary with mating success.. We also carry out field behavioral assays to test whether male pheromones serve as long range attractants for both males and females.

## METHODS AND MATERIALS

### Sampling

Male bees were collected between April 2020 and May 2022 while actively displaying in the late afternoon. Samples were collected throughout the city of Davis, California, USA. For a detailed list of sites, see supplementary materials (Table S1). In addition, two males were collected by hand from their nest in a log prior to the commencement of their mating season. These latter samples were excluded from the overall analysis as they were collected using a different method and later found to lack the main pheromone compounds. Upon collection, the samples were frozen at at -20°C. First, we measured the intertegular distance, which we use as a proxy for body size (Cane 1987). We also recorded the wing wear (nicked wing edges), which we used as a proxy for male age (Mueller and Wolf-Mueller 1993). To dissect the thoracic pheromone gland from the mesothorax, the recently frozen bees were pinned onto an agar dissection dish, with one pin placed through the prothorax and an additional pin placed through the abdomen. Using forceps, the dorsal mesosomal cuticle was cut away, exposing the mesosomal gland below. The glands were removed from the thorax, placed onto a white microscope stage, and photographed, to measure their color. The camera, a Cannon EOS 60D, was mounted on a tripod, approximately 40 centimeters above the microscope stage, and manually focused during each sample. Lighting was provided using a portable microscope lamp with two bulbs, placed on opposing sides of the stage, and diffused through wax paper.

### Coloration Analysis

The Fiji distribution of ImageJ was used to collect color data (Schindelin et al. 2012). In each photo, the gland was hand selected, five random points were chosen across the gland, and across these five points, the program provided both the RGB (red-green-blue color model) values and the HSB (hue-saturation-brightness color model) values for the image. These values were then averaged. To assess color, a ratio of R (red) and G (green) was calculated, as the different color values are only meaningful relative to the other values, and this ratio was used in our analysis (Bergman and Beehner 2008).

### Chemical Analysis

Each thoracic gland was placed in a glass vial containing 500μl hexane with 5μg of 2-undecanone as an internal standard and solvent extracted for one hour. Following this, the solvent was transferred to a new vial, and stored at -20°C. Samples were analyzed using an Agilent model 5977A mass-selective detector connected to Agilent GC model 7890B, with a HP-5 Ultra Inert column (Agilent, 117 30 m × 0.25 mm, 0.25 µm). 1μl of each sample was injected using Agilent ALS 7694 autosampler in split mode with a 5:1 ratio with helium as the carrier gas (250°C injector temperature, split flow of 3.5 ml/min). The temperature program started at 80°C for 3 minutes, and then rose at 10°C/min to 300°C. The temperature was held at 300°C for 3 minutes and 315°C for 3 minutes. We quantified components using an internal standard of 2-undecanone, and compounds not found in at least 10% of individuals were disregarded. Compounds were identified by comparing to prior work done on this system (Andersen et al. 1988) and to NIST reference libraries. Furthermore, we identified compounds all trans-geranylgeraniol and all trans-farnesal using authentic standards purchased from Sigma-Aldrich chemicals.

### Behavioral Assay

We conducted field behavioral bioassays in the spring of 2023 at a pair of test sites in the UC Davis Arboretum (38.5328, -121.7547). The sites were 10 meters apart, on the same species of shrubby plant. On each study day, a coin was flipped to randomly assign one of two treatments to each site. During the study, each site had a qualitative cellulose filter paper disc (Sterlitech, Catalog # CFP3-090), with a diameter of 9 cm, suspended from a branch of the shrub approximately 1.5 meters off the ground using an odorless, green lab tape. To create the experimental mixture, we used a mixture of the actual extracts from the top 75% of males in terms of total pheromone abundance (n=30) from the samples included in this study. We applied 1ml of this mixture to the filter paper in the field. Since each male’s gland was extracted into 0.5 ml of solvent, each 1ml aliquot represents two males’ worth of pheromone. The second treatment was a control, which consisted of 1 ml of pure hexane, also applied to the filter paper discs in the field.

Following the application of the two treatments, the site was observed for the following 60 minutes and all interactions from both male and female bees were recorded. Interactions were counted if an individual altered their flight path and approached the treated papers within 1 meter. We also noted if a “strong” interaction occurred, in which case the individual stopped and hovered within 0.5 meter of the filter paper for any length of time. We chose to observe changes in flight path and hovering as these are likely to be the initial stages of mate finding. Mating attempts would not be expected with solely chemical cues because visual cues are also required in this system for mating (Marshall and Alcock 1981). Initially, observations took place from 16:00-17:00 in the afternoon, coinciding with male *X. sonorina* activity. However, throughout the season, this shifted back to 16:30-17:30 and eventually 17:00-18:00, following the observed male display behavior, which progressed later into the afternoon as the season advanced (Table S7). We carried out 14 assays between mid-May and the end of June 2023. Previous observations found that male activity only happens at higher temperatures (>26.5°C). We, therefore, only carried out the assay on days with temperatures high enough to observe males displaying on nearby plants.

### Statistical Analysis: Variation in Compound Amount

Firstly, we were interested in determining the factors correlated with variation in total compound amount produced by the thoracic gland. We tested the effects of intertegular distance, wing wear, and various characteristics of the glands appearance, including color (R:G ratio), hue, saturation, and brightness, on total compound abundance, using ANOVA. We then evaluated our model fit with Akaike’s information criteria (AIC) using the *AICcmodavg* package (Mazerolle 2020). The model with the lowest AIC score was chosen as the final model, containing the most likely explanatory variables.

### Statistical Analysis: Variation in Chemical Profile Composition

The univariate analysis only includes total compound amount and does not consider which compounds are present in the chemical profile. To include this information in the analysis, we carried out a multivariate analysis to analyze the chemical mixture of the males, taking all the compounds into account. Due to the zeros present in our data matrix, we visualized divergence across the samples using nonmetric multidimensional scaling (NMDS) in three dimensions, based on a Bray–Curtis similarity matrix created using absolute peak areas. We created the NMDS with the “metaMDS” function in *vegan,* and this was plotted using the *ade4* package (Thioulouse et al. 2018; Oksanen et al. 2022).

To investigate factors that explain pheromone composition, we used a PERMANOVA (permutational multivariate analysis of variance) on a Bray-Curtis similarity matrix, using the “adonis2” function in *vegan* with 1000 permutations (Oksanen et al. 2022). We assessed each term individually using the option by=”margin”, this ensures that the order of terms in the model does not affect the results (in contrast to the by=”sequential” option where each term is considered in order). Model fit was evaluated using AIC scores, with each model being tested sequentially. The model with the lowest AIC score was selected as the final model and identified as the model containing the explanatory factors. However, we chose the simplest model as the best fit if we found that two models were within two AIC values of each other. We also repeated this analysis with the relative compound abundances.

### Statistical Analysis: Variation in the Most Abundant Compounds

The chemical profile of the bees is dominated by three main compounds, as described previously (Andersen et al. 1988). These compounds appear to be biologically important to the reproduction of *X. sonorina*, since an artificial mixture of two of these compounds, all trans-farnesal and all trans-geranylgeraniol, seems to be sufficient to attract females (Minckley et al. 1991). Therefore, we re-analyzed our data including just these three primary compounds to determine which factors are correlated with their abundance and composition as described for the analysis of the entire dataset. We also carried out Pearson’s correlation tests for these compounds with both size and brightness.

### Plotting and Data Manipulation

To make plots we used the following packages: *ggpubr* (Kassambara 2020), *cowplot* (Wilke 2020) and *ggplot2* (Wickham 2016). In addition we used *tidyr* (Wickham and Girlich 2022), *plyr* (Wickham 2011), and *readr* (Wickham et al. 2023) for data transformation. Analyses were carried out in R version 4.2.1 (R Core Team 2022).

### Behavioral Assay Analysis

To determine if the likelihood of interaction was related to either sex of the individuals or treatment (male gland extract vs. hexane control), we used a binomial GLM (generalized linear model), with the package *MASS* (Venables and Ripley 2002). The response variable was whether at least one interaction was observed in a given trial. We included sex and treatment as explanatory variables as well as an interaction term between the two. To test significance of factors, we used likelihood ratio tests on models with and without the variable of interest.

All raw data and analyses are available on the OSF (https://osf.io/uf87d/?view_only=ab07af2f3c4e4c05a5f9f5fad680dd64).

## RESULTS

We sampled a total of 40 males and identified 43 different compounds in their chemical profile. On average, 284.5±168.7 μg (SD) of total compound were present in each sample.

Across all the samples, the three main compounds described by prior studies in an Arizona population were also found in the Davis population, with all trans-geranylgeraniol, all trans-farnesal, and 3,7,11-trimethyl-2,7,10-dodecatrienal being the three most abundant compounds (Figure 1, Panel B, labels 1, 2, 3) (Andersen et al. 1988).

In the sampled displaying males, we observed a wide range of gland color, with some much darker than others (Figure S1). In addition to the displaying males, we also sampled two adult males which were collected from their nest in a log prior to the start of the mating season. Upon dissection, we observed that these males had extremely pale, almost white mesosomal glands, and when chemically analyzed, these males had an average total compound amount of 2.08±1.1 μg (SD). Additionally, these males lacked the three main compounds seen in the actively displaying males.

### Variation in Compound Amount

To investigate the impact of different factors on total compound amount, we carried out a univariate ANOVA analysis. We found that both brightness and size were significantly related to the total amount of compound (Brightness: ANOVA, F_1,36_= 10.99, p=0.002094; Size: ANOVA, F_1,36_=14.02, p=0.000632) (Figure 2, Table S2). We found that males with brighter glands have an overall lower total compound amountThese results agree with similar findings about the glands progressing through a pre-secretory (lightest = brightest color) phase, then into an active secretor phase, and finally a dark reddish brown storage phase, at which point males will have accumulated their maximum store of pheromones (Buchmann, unpubl.). We also found that smaller males, measured by their intertegular distance, produce a lower total compound amount. We did not find that wing wear significantly explained variation in compound amount.

**Figure 2:**
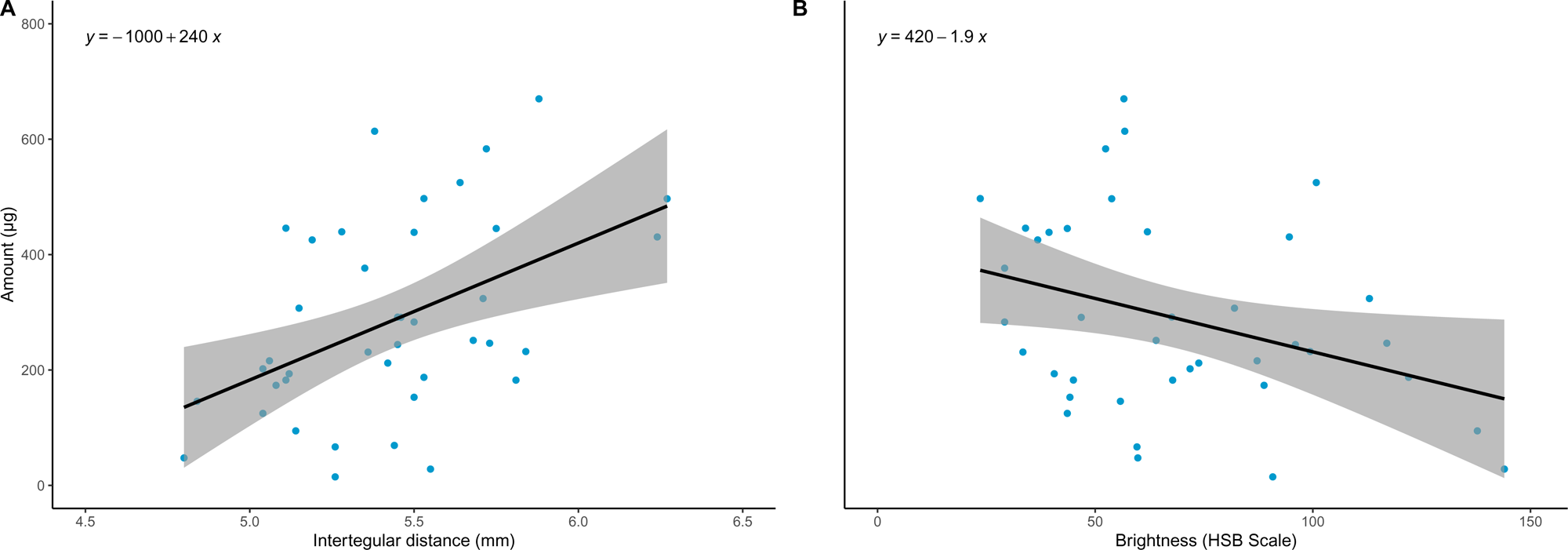
(a) Amount of pheromone is positively related to male *X. sonorina* body size (ANOVA, F_1,36_=14.02, p=0.000632), using intertegular distance as a proxy. (b) Amount of pheromone is negatively related with gland brightness (ANOVA, F_1,36_= 11.32, p=0.001832), according to the HSB scale. A higher value indicates a lighter color of the gland.

### Variation in Chemical Profile Composition

To identify which factors drive overall compositional differences in the chemical profile of male glands, we carried out a multivariate PERMANOVA analysis. As in the univariate analysis, the best model included both brightness (PERMANOVA, Brightness, *F*1,36 = 5.1835, *p* =0.001998) and size (PERMANOVA, Size, *F*1,36 = 7.5555, *p* =0.000999) (Figure 3, Table S3). We found that size accounts for 16% of observed variation in chemical profile, with a further 11% explained by gland brightness. We did not find that wing wear significantly explained variation in chemical profile.

**Figure 3:**
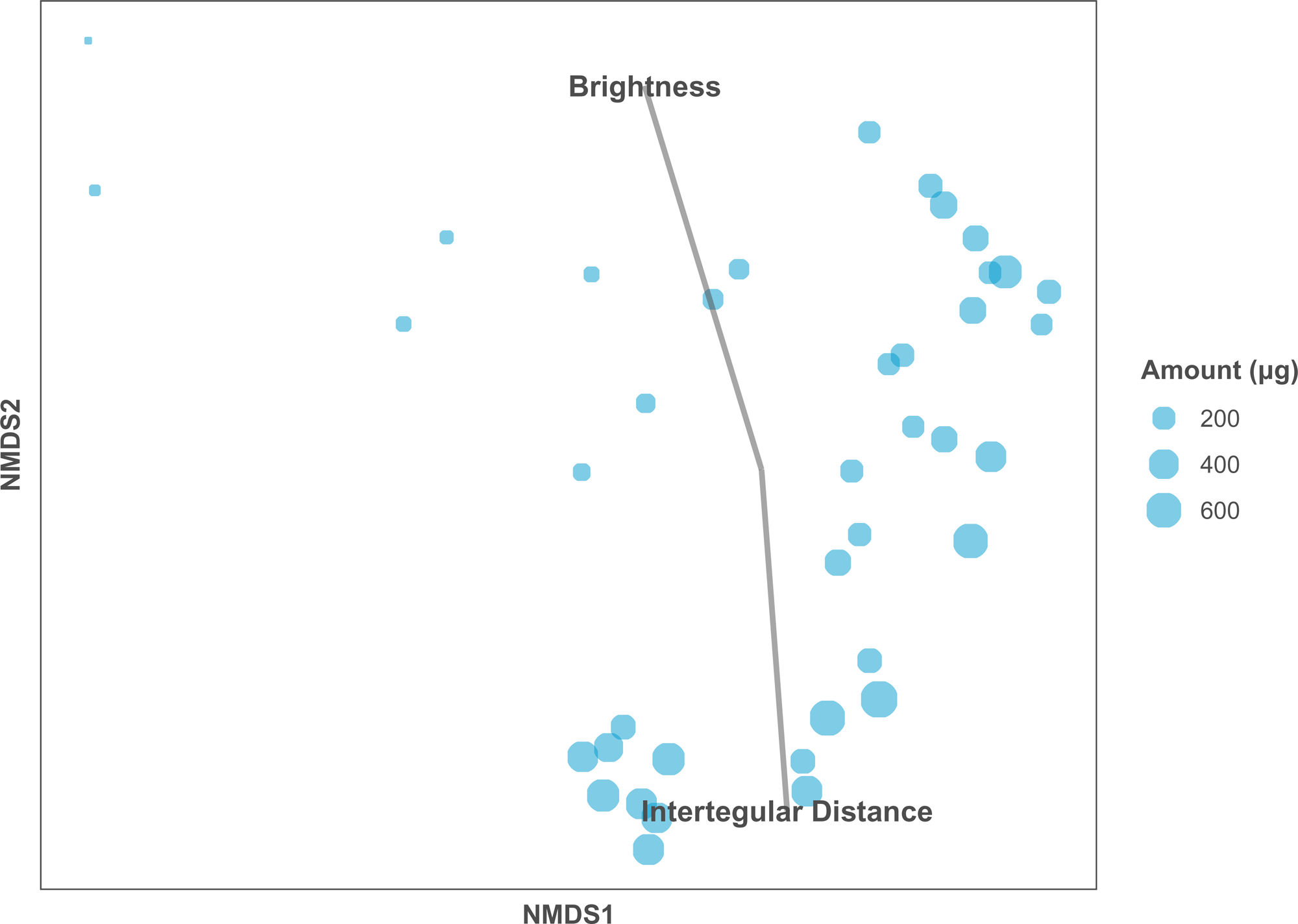
NMDS (nonmetric multidimensional scaling) plot illustrating in two dimensions the variation in mesosomal gland contents of male *X. sonorina.* Size of each circle corresponds to the absolute amount of pheromone. The impact of body size (with intertegular distance as a proxy) and brightness are shown.

When we reanalyzed the data using relative compounds abundances, we found that none of our variables explained variation in chemical profile (Table S4).

### Variation in the Most Abundant Compounds

The prior two analyses included every compound which was present in at least 10% of sampled males. However, prior studies have identified three main compounds, two of which can attract females, and so we re-ran the previous analyses on a reduced dataset of these three main compounds (Andersen et al. 1988; Minckley et al. 1991). Combined, these three main compounds made up an average of 250.9 ± 157.7 μg (SD) in each male gland, which is 86.4% ± 6.02% (SD) of total gland pheromone amount.

We found that both brightness and size was correlated with the total amount of compound (Brightness: ANOVA, F(_1,36_)= 11.32, p=0.001832; Size: ANOVA, F(_1,36_)=13.66, p=0.000724)(Figure S2, Table S5). To identify which factors drive overall compositional differences in the three main compounds, we carried out a PERMANOVA analysis. The best model fit again included both brightness (PERMANOVA, Brightness, *F*1,36 = 5.1614, *p*=0.003996) and male bee size (PERMANOVA, Size, *F*1,36 = 7.9076, *p* =0.000999), with size explaining 17% of observed variation in the three main compounds, and brightness explaining a further 11% (Figure S3, Table S6). We did not find that wing wear significantly explained variation in the three main compounds. These results were coincident with the entire chemical blend, suggesting that these compounds are responsible for the patterns seen when analyzing the entire dataset.

To determine if all three of these compounds are correlated with brightness and size, we analyzed each compound individually. We found that variation in both all trans-farnesal and all trans-geranylgeraniol is related to size and brightness (all trans-farnesal: Size, ANOVA, F(_1,36_)= 9.1, p=0.0047; Brightness, ANOVA, F(_1,36_)= 10.9, p=0.0022)(all trans-geranylgeraniol: Size, ANOVA, F(_1,36_)= 11.88, p=0.0015; Brightness, ANOVA, F(_1,36_)= 7.72, p=0.0086). We did not find that variation in 3,7,11-trimethyl-2,7,10-dodecatrienal was explained by size or brightness. We also calculated the strength of correlation between all trans-farnesal and all trans-geranylgeraniol with size and brightness. We found that all trans-farnesal is significantly correlated with both size (Pearson’s correlation, r=0.4, p=0.011) and brightness (Pearson’s correlation, r=-0.37, p=0.019). In contrast, all trans-geranylgeraniol is only significantly correlated with size (Pearson’s correlation, r=0.45, p=0.0032), but not brightness (Pearson’s correlation, r=-0.3, p=0.068). Taken together, these two compounds are responsible for the main patterns detected in the overall dataset (Figure 4).

**Figure 4:**
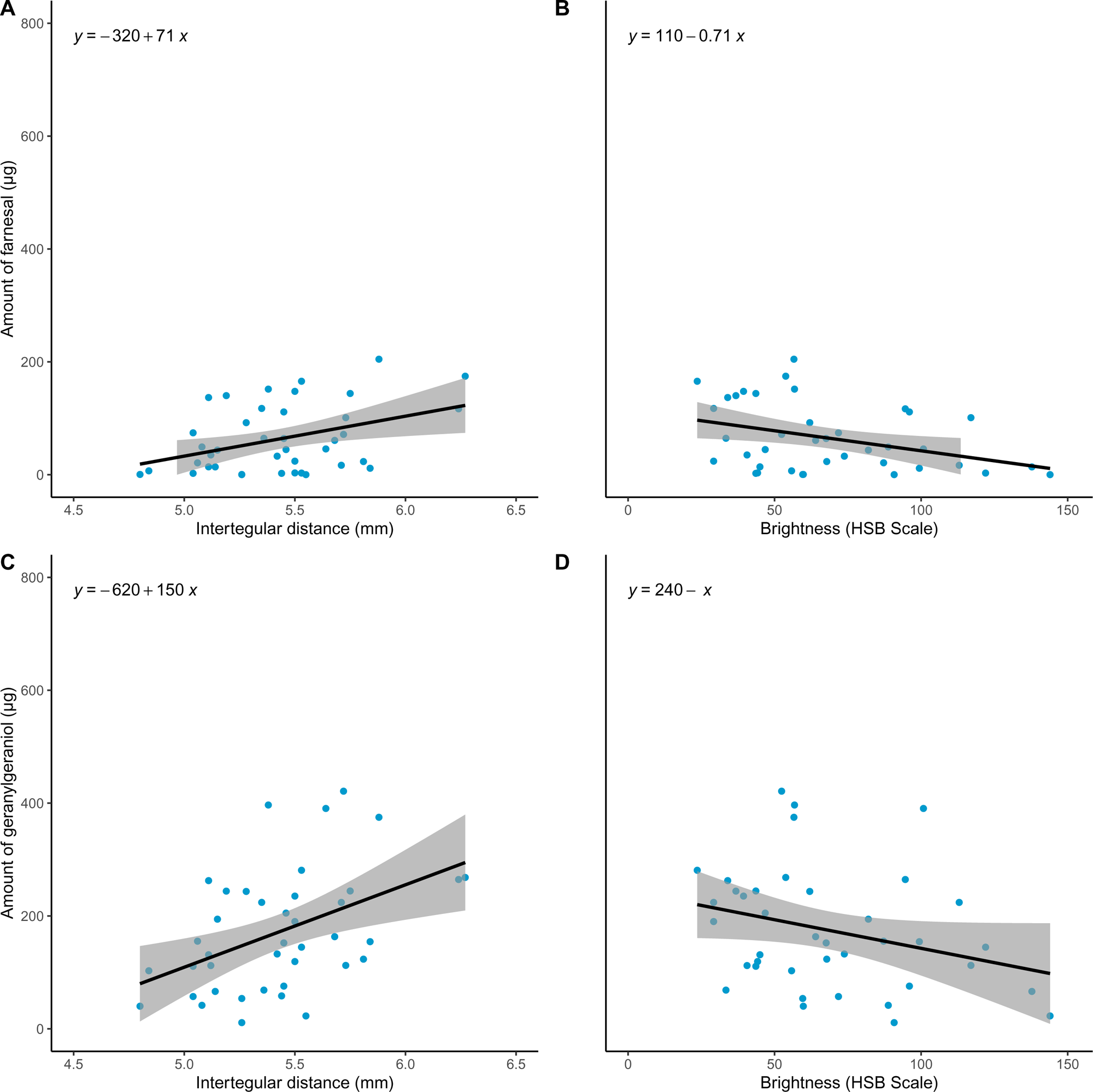
(a) Scatterplot of the relationship between male *X. sonorina* body size, with the intertegular distance as a proxy, and the absolute abundance of all trans-farnesal. (b) Scatterplot of the relationship between the absolute abundance of all trans-farnesal and brightness of the mesothoracic gland, measured on the HSB scale, with a higher value corresponding to a lighter colored gland. (c) Scatterplot of the relationship between male *X. sonorina* body size, with the intertegular distance as a proxy, and the absolute abundance of all trans-geranylgeraniol. (d) Scatterplot of the relationship between the absolute abundance of all trans-geranylgeraniol and brightness of the mesothoracic gland, measured on the HSB scale, with a higher value corresponding to a lighter colored gland.

### Behavioral Assay Results

In total, we conducted 14 behavioral field trials (bioassays), and interactions by either males or females occurred in 13 of the 14 trials, resulting in 37 total interactions (Table 1, Table S7). Females interacted with the control treatment on 4/14 days, while males interacted with the control on 1/14 days. These numbers were much higher for the filter paper treated with extract, with females interacting 11/14 days, and males 8/14 days. Treatment was found to significantly explain the likelihood of interaction (2ΔlnL= 16.016, d.f.=1, p=0.000063).

We also observed what we have termed “strong responses”. These consist of individuals stopping and hovering within 0.5 meters of the filter paper. This occurred on 3 separate days for the males, and 5 separate days for the females, and was observed solely on the filter paper with the male’s extract applied, while this hovering behavior did not occur towards the control (Table S7).

## DISCUSSION

*Xylocopa sonorina* males release large amounts of their sex pheromone during mating displays which are detectable to humans as a rose-like scent (Alcock and Smith 1987). Here, we investigated the chemical ecology of *X. sonorina*, describing pheromone variation within a population in California and identified morphological factors which could explain the observed variation. As previously identified in Arizona individuals, we found that the three main constituents of this gland were all trans-geranylgeraniol, all trans-farnesal, and an isomer of 3,7,11-trimethyl-2,7,10-dodecatrienal (Andersen et al. 1988). This shows a high level of consistency and stability in this chemical signal across the range of this species, and in two areas with different climates. We show that larger males with darker gland coloration (*i.e.* mature glands) have more total pheromone in their glands. We found that this is mainly by variation in the abundance of all trans-farnesal and all-trans geranylgeraniol, two of the major compounds found in the gland. These two compounds have also previously been shown to be sufficient to attract females (Minckley et al. 1991), a result which we further support here, and provide new evidence that males are also attracted to male pheromones.

We found that male size explained the most observed variation in male pheromone amount in *X. sonorina*. In bees, as with other holometabolous insects, the male’s adult body size is determined during the larval stage, and is heavily impacted by both abiotic (temperature, latitude, altitude, season) and biotic (primarily diet) factors (Chole et al. 2019). Larger male body size is thought to signal a higher quality diet during development, which corresponds to increased mating success (Agosta 2010). We found that larger *X. sonorina* produce more sex pheromone, which we hypothesize could be related to a higher mating success, as seen across a range of insects. For example, in the tobacco moth, *Ephestia elutella*, larger males produce more pheromone when compared to smaller males, and these pairings result in larger offspring, with size being positively correlated with mating success in these moths (Phelan and Baker 1986). We hypothesize that larger male *X. sonorina* which produce a greater abundance of their sex pheromones could allow these larger males to attract females from a greater distance than smaller males, or potentially to release pheromones for a longer period of time. However, further work is required to discern if females show a preference for these larger males.

We also found that male gland color was correlated with pheromone amount. In some cases, age could be signaled by sex pheromones, ensuring that mating occurs in the window between sexual maturation and senescence (Gomez-Diaz, Carolina and Benton, Richard 2013). In the butterfly *Bicyclus anynana*, the male’s sex pheromone composition changes as they age, and females exhibit preferences for the pheromones of middle aged males (Nieberding et al. 2012), with similar patterns documented in other species (Lassance and Löfstedt 2009; Chemnitz et al. 2015). We did not find a relationship between male *X. sonorina* sex pheromone and age when using wing wear as a proxy; however this may be a poor indicator of age due to variability in wing degradation (Eltz et al. 1999). We decided to use the color of the gland as a proxy for male maturation as previous work on *X. sonorina* has found the gland is initially light in color and subsequently darkens into a sclerotized storage organ with many hair lined tubules (Buchmann pers. obs.; Minckley et al. 1991; Ostwald et al. 2022). During our dissections, we observed a range of colors of the mesosomal glands, from near white in some that were collected prior to the commencement of the males display season to very dark red-brown in some displaying males. The two males collected from their nests before their display season began had nearly white mesosomal gland tissues containing only trace amounts of pheromone. We found that the brightness of the gland correlates with pheromone amount, with darker glands containing higher amounts of pheromone, in agreement with the prediction that older males have higher pheromone amounts.

The effect of higher sex pheromone production by larger, more sexually mature male *X. sonorina* on female choice is unclear. Two ways in which sex pheromone production could impact female choice are through the effectiveness of the pheromone as a long-range attractant or as an indicator of mate quality. Based on our behavioral assays, we present preliminary evidence that these compounds serve as a long-range attractant for both males and females. In hymenopterans, it has long been accepted that sex pheromones serve as important facilitators of sexual reproduction by acting as long-range attractants (Ayasse et al. 2001). Within *X. sonorina,* an early study showed that the primary constituents from the male gland seemed to be sufficient to attract females (Minckley et al. 1991). The results from our behavioral assay provide further evidence to support those observations.

Nonetheless, this does not discount the possibility that females are assessing male quality through their sex pheromones. *Xylocopa sonorina* exhibits a non-resource-based mating system. This is in contrast to the mating system of two other *Xylocopa* species found in California, *X. tabaniformis* and *X. californica.* In these species, males defend resource-based territories, have enlarged eyes and lack the large mesosomal glands found in *X. sonorina.* This suggests that as *Xylocopa* bees shifted from a resource based, territorial mating strategy to a non-resource-based mating strategy, they shifted from a visual based mating system to one that is facilitated by pheromones. In the *X. sonorina* system, females must land for mating to commence (Marshall and Alcock 1981), and mating occurs relatively infrequently, with only 24% of female interactions with displaying males resulting in copulation (Alcock and Johnson 1990). It appears that females are assessing multiple males, often in a lekking display, with status communicated through chemical signals. However, further studies are required to test this hypothesis and determine the effect that the male’s mesosomal glands play in the female assessment.

As well as being used by females to locate and assess mate quality, sex pheromones may also be used to facilitate male-male interactions. When conspecific, displaying *X. sonorina* males encounter each other at the same display location, they may exhibit one of two different behavioral responses. Sometimes, one male will emerge dominant and chase off their competitor, while in other cases, up to several males may share a display site, flying in loose circles around each other and the focal point for some length of time (Alcock and Smith 1987). Further studies are needed to determine if this variation in behavior is reflected in the chemistry of the pheromones released by displaying male *X. sonorina*. Based on results from our behavioral assay, we provide evidence that male *X. sonorina* are attracted to the mesosomal chemical blend released by males during their mating display. Additionally, similar to what has been described in other hymenopterans, such as sphecid (Coelho and Holliday 2001) and pompilid wasps (Alcock and Kemp 2006), larger males may be better fliers or a have a better ability to defend their territory. *Xylocopa sonorina* may provide a viable system to investigate how sex pheromones facilitate male-male interactions during their complex mating displays.

Our study reveals variation in sex pheromone amounts are correlated with both body size and gland color in male *X. sonorina*. Larger, more sexually mature males produce higher amounts of sex pheromone. In addition, we show preliminary evidence that this blend of male pheromones serves as a long-range attractant to both the opposite and same sex. Future studies could clarify the fitness consequences of variation in pheromone amount, whether by impacting attraction range, female choice, or male-male competitive interactions.

## Acknowledgements

We acknowledge the support and feedback of the Ramirez lab with this project.

## STATEMENTS AND DECLARATIONS

### Author Contributions

All authors contributed to the study conception and design. Material preparation, data collection were performed by Andrew J. Goffinet, Kathy Darragh, and Nicholas Saleh. The analysis was carried out by Andrew J. Goffinet and Kathy Darragh. The first draft of the manuscript was written by Andrew J. Goffinet, and all authors commented on previous versions of the manuscript. All authors read and approved the final manuscript.

