## Supplemental Materials for "Individual variation in male pheromone production in *Xylocopa sonorina* correlates with size and gland color"

### Individual variation in male pheromone production in carpenter bees correlates with size and sexual maturity.

Journal of Chemical Ecology

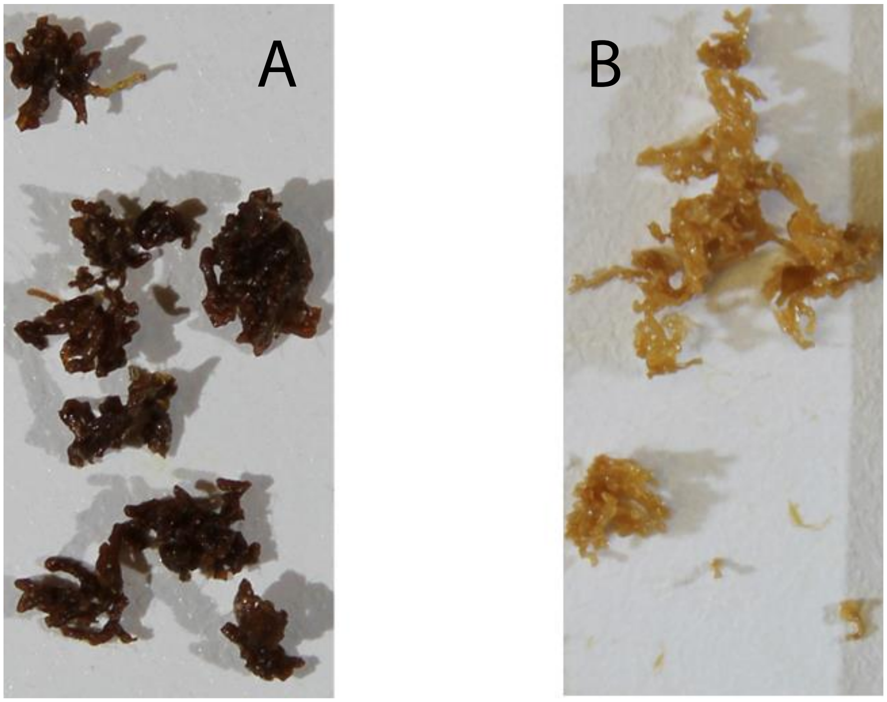


**<u>Figure S1:</u>** (A) Photo of a dark gland from a displaying male (X0007) which contains 182.5 $\mu$g of pheromone. (B) Photo of a lighter gland from a displaying male (X0020) which contains 28.2$\mu$g of pheromone.

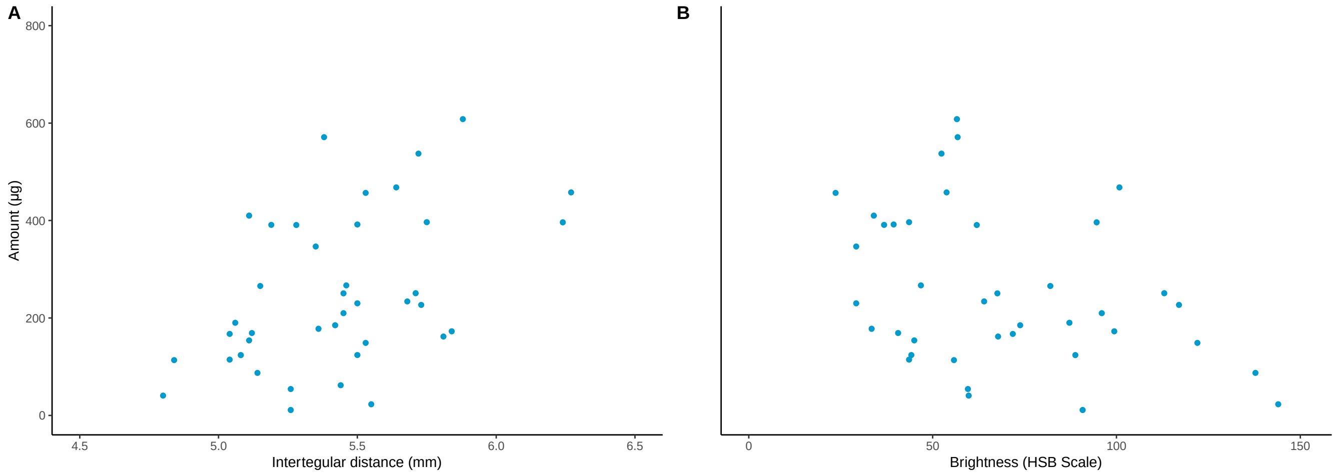


**<u>Figure S2:</u>** (a) Scatterplot of the relationship between male *X. sonorina* body size, with the intertegular distance as a proxy, and the absolute abundance of the three primary constituents. (b) Scatterplot of the relationship between the absolute abundance of the three primary compounds and brightness of the mesothoracic gland, measured on the HSB scale, with a higher value corresponding to a lighter colored gland.

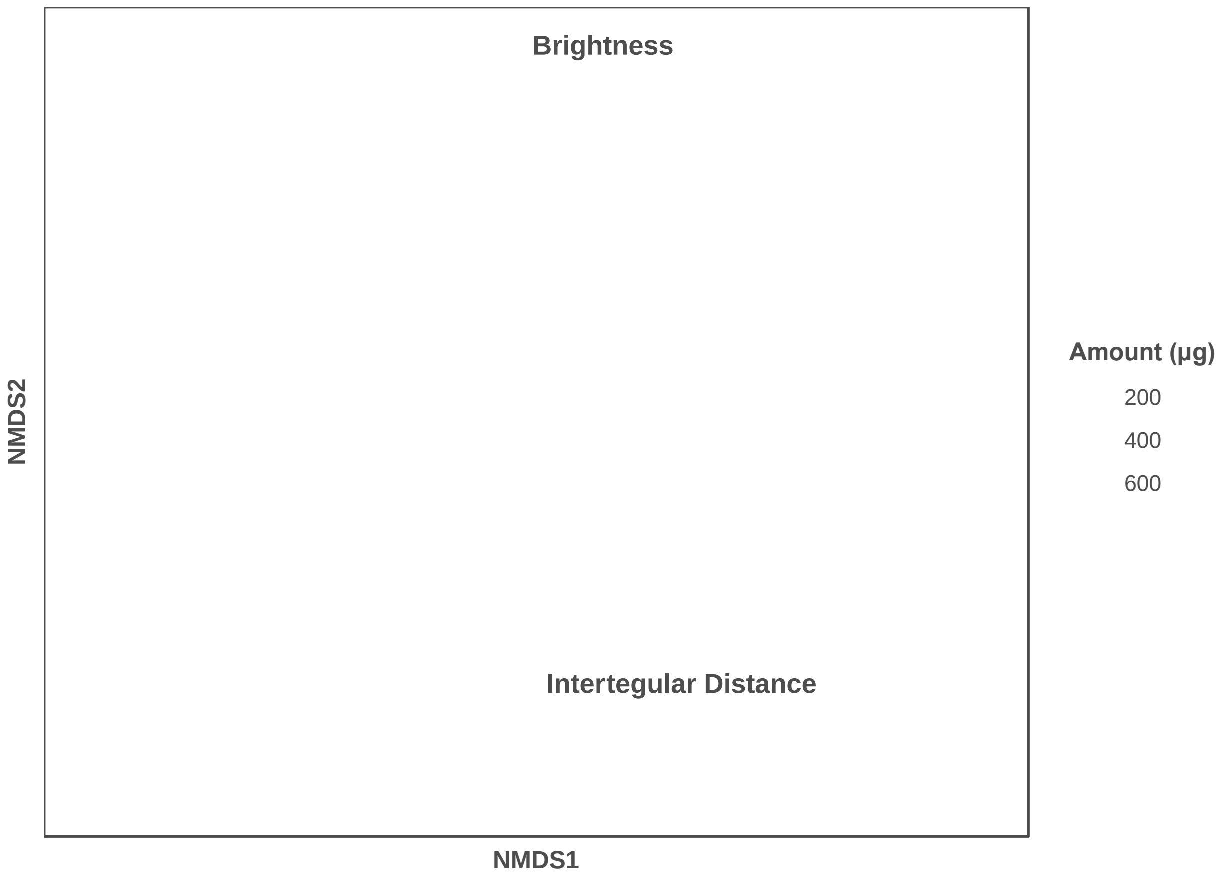


**Figure S3:** NMDS (nonmetric multidimensional scaling) showing in two dimensions the variation in the chemical profile of the three primary compounds extracted from male *X. sonorina* mesothoracic glands. The size of each circle corresponds to the absolute amount of these three compounds, while the factors of body size (with intertegular distance as a proxy) and brightness are labeled.

**Table S1: Information on sample locations**

| State | County | Latitude | Longitude | Number of Samples | Site Description |
|---|---|---|---|---|---|
| California | Yolo/Solano | 38.55 | -121.75 | 11 | Town of Davis (location not specified) |
| California | Yolo | 38.5411 | -121.7552 | 1 | Bushes near SE corner of botanical conservatory on UCD campus |
| California | Yolo | 38.570 | -121.764 | 9 | Bushes on northern and southern edge of the Northstar Park pond |
| California | Yolo | 38.551 | -121.769 | 2 | Westwood townhouse Apartments |
| California | Yolo | 38.554 | -121.774 | 6 | Arroyo Park |
| California | Yolo | 38.5420 | -121.7557 | 1 | BOG Garden north of Storer Hall |
| California | Yolo | 38.551 | -121.773 | 3 | Westwood Park |
| California | Yolo | 38.545 | -121.745 | 2 | Central Park |
| California | Yolo | 38.537 | -121.748 | 6 | Shrubs and trees along the UC Davis Arboretum |

| | | | | | |
|---|---|---|---|---|---|
| California | Yolo | 38.555 | -121.766 | 1 | Sycamore Park |

**Table S2: Model selection for anova analysis of total compound abundance**

| Model | Residual Sum of Squares | DF | AIC |
|---|---|---|---|
| Total Compound Amount ~ Size + Color (R:G) + Saturation + Brightness + Hue + Wingwear | 577399 | 32 | 505.98 |
| Total Compound Amount ~ Size + Color (R:G) + Saturation + Brightness + Wingwear | 579021 | 33 | 502.91 |
| Total Compound Amount ~ Size + Saturation + Brightness + Wingwear | 592781 | 34 | 500.83 |
| Total Compound | 600189 | 35 | 498.51 |

| | | | |
|---|---|---|---|
| Amount ~ Size + Wingwear + Brightness | | | |
| **Total Compound Amount ~ Size + Brightness** | 626711 | 36 | 497.56 |
| Total Compound Amount ~ Size * Wingwear * Brightness | 579487 | 31 | 509.53 |

Note- If two models were within 2 AIC points of each other, we selected the simpler model as the most parsimonious. The selected model is bolded.

**Table S3: Model selection for PERMANOVA analysis of chemical profiles using absolute compound abundances**

| Model | Residual Sum of Squares | DF | AIC |
|---|---|---|---|
| Chemical Profile ~ Color (R:G) + Hue + Saturation + Brightness + Size + Wingwear | 3.0324 | 32 | 57.2644 |
| Chemical Profile ~ Color (R:G) + Hue + | 3.0937 | 33 | 56.04491 |

| | | | |
|---|---|---|---|
| Saturation + Brightness + Size | | | |
| Chemical Profile ~ Hue + Saturation + Brightness + Size | 3.1624 | 34 | 54.90172 |
| Chemical Profile ~ Saturation + Brightness + Size | 3.2839 | 35 | 54.37213 |
| **Chemical Profile ~ Brightness + Size** | 3.3814 | 36 | 53.51381 |
| Chemical Profile ~ Saturation * Brightness * Size | 3.0120 | 31 | 59.00168 |

Note- If two models were within 2 AIC points of each other, we selected the simpler model as the most parsimonious. The selected model is bolded.

**Table S4: Model selection for PERMANOVA analysis of chemical profiles using relative compound abundances**

| Model | Residual Sum of Squares | DF | AIC |
|---|---|---|---|
| Chemical Profile ~ Color (R:G) + Hue + Saturation + Brightness + Size + | 1.24298 | 32 | 22.483 |

| | | | |
|---|---|---|---|
| Wingwear | | | |
| Chemical Profile ~ Color (R:G) + Saturation + Brightness + Size + Wingwear | 1.23909 | 33 | 20.36062 |
| Chemical Profile ~ Color (R:G) + Brightness + Size + Wingwear | 1.25003 | 34 | 18.70358 |
| Chemical Profile ~ Color (R:G) + Brightness + Size | 1.26527 | 35 | 17.17603 |
| Chemical Profile ~ Color (R:G) + Brightness | 1.31265 | 36 | 16.60996 |
| Chemical Profile ~ Color (R:G) | 1.35276 | 37 | 15.7836 |

Note- If two models were within 2 AIC points of each other, we selected the simpler model as the most parsimonious. No model resulted in significant explanatory factors.

**Table S5: Model selection for anova analysis of three main compounds**

| Model | Residual Sum of | DF | AIC |
|---|---|---|---|

| | | | |
|---|---|---|---|
| | Squares | | |
| Total Compound Amount ~ Size + Color (R:G) (R/G) + Saturation + Brightness + Hue + Wingwear | 506799 | 32 | 500.90 |
| Total Compound Amount ~ Size + Color (R:G) + Saturation + Brightness + Wingwear | 508687 | 33 | 497.86 |
| Total Compound Amount ~ Size + Saturation + Brightness + Wingwear | 518253 | 34 | 495.59 |
| Total Compound Amount ~ Size + Wingwear + Brightness | 524110 | 35 | 493.23 |
| **Total Compound Amount ~ Size + Brightness** | 550794 | 36 | 492.52 |

| | | | |
|---|---|---|---|
| Total Compound Amount ~ Size * Wingwear * Brightness | 506294 | 31 | 504.27 |

Note- If two models were within 2 AIC points of each other, we selected the simpler model as the most parsimonious. The selected model is bolded.

**Table S6: Model selection for PERMANOVA analysis of three main compounds**

| Model | Residual Sum of Squares | DF | AIC |
|---|---|---|---|
| Chemical Profile ~ Color (R:G) + Hue + Saturation + Brightness + Size + Wingwear | 3.0239 | 32 | 57.15487 |
| Chemical Profile ~ Color (R:G) + Hue + Saturation + Brightness + Size | 3.0891 | 33 | 55.98688 |
| Chemical Profile ~ Hue + Saturation + Brightness + Size | 3.1789 | 34 | 55.10506 |
| Chemical Profile ~ Saturation + | 3.2893 | 35 | 54.4368 |

| | | | |
|---|---|---|---|
| Brightness + Size | | | |
| **Chemical Profile ~ Brightness + Size** | 3.3921 | 36 | 53.63708 |
| Chemical Profile ~ Saturation * Brightness * Size | 3.0280 | 31 | 59.20782 |

Note- If two models were within 2 AIC points of each other, we selected the simpler model as the most parsimonious. The selected model is bolded.

**Table S7: Raw data from behavioral assay**

| Trial Number | Treatment | Trial Date | Start Time | Number Male Interactions | Number Female Interactions |
|---|---|---|---|---|---|
| 1 | Extract | 16 May 2023 | 16:00 | 4 (1) | 3 (1) |
| 1 | Control | 16 May 2023 | 16:00 | 0 | 2 |
| 2 | Extract | 17 May 2023 | 16:30 | 0 | 1 |
| 2 | Control | 17 May 2023 | 16:30 | 0 | 1 |
| 3 | Extract | 19 May 2023 | 16:30 | 0 | 1 (1) |
| 3 | Control | 19 May 2023 | 16:30 | 2 | 1 |

| | | | | | |
|---|---|---|---|---|---|
| 4 | Extract | 21 May 2023 | 17:00 | 1 (1) | 1 (1) |
| 4 | Control | 21 May 2023 | 17:00 | 0 | 0 |
| 5 | Extract | 22 May 2023 | 16:30 | 0 | 2 |
| 5 | Control | 22 May 2023 | 16:30 | 0 | 1 |
| 6 | Extract | 31 May 2023 | 16:30 | 0 | 3 (1) |
| 6 | Control | 31 May 2023 | 16:30 | 0 | 0 |
| 7 | Extract | 13 June 2023 | 16:30 | 1 | 2 |
| 7 | Control | 13 June 2023 | 16:30 | 0 | 0 |
| 8 | Extract | 14 June 2023 | 16:30 | 1 (1) | 1 (1) |
| 8 | Control | 14 June 2023 | 16:30 | 0 | 0 |
| 9 | Extract | 20 June 2023 | 17:00 | 2 | 0 |
| 9 | Control | 20 June 2023 | 17:00 | 0 | 0 |
| 10 | Extract | 21 June 2023 | 17:00 | 1 | 1 |
| 10 | Control | 21 June 2023 | 17:00 | 0 | 0 |
| 11 | Extract | 26 June 2023 | 17:00 | 1 | 1 |
| 11 | Control | 26 June 2023 | 17:00 | 0 | 0 |

| | | | | | |
|---|---|---|---|---|---|
| 12 | Extract | 27 June 2023 | 17:00 | 1 | 0 |
| 12 | Control | 27 June 2023 | 17:00 | 0 | 0 |
| 13 | Extract | 28 June 2023 | 17:00 | 0 | 2 |
| 13 | Control | 28 June 2023 | 17:00 | 0 | 0 |
| 14 | Extract | 23 May 2023 | 16:30 | 0 | 0 |
| 14 | Control | 23 May 2023 | 16:30 | 0 | 0 |

Note: The number in parentheses are “strong” interactions observed.
